# Segmentation of Pancreatic Ductal Adenocarcinoma (PDAC) and surrounding vessels in CT images using deep convolutional neural networks and texture descriptors

**DOI:** 10.1101/2021.06.09.447508

**Authors:** Tahereh Mahmoudi, Zahra Mousavi Kouzahkanan, Amir Reza Radmard, Raheleh Kafieh, Aneseh Salehnia, Amir H. Davarpanah, Hossein Arabalibeik, Alireza Ahmadian

## Abstract

Fully automated and volumetric segmentation of critical tumors may play a crucial role in diagnosis and surgical planning. One of the most challenging tumor segmentation tasks is localization of Pancreatic Ductal Adenocarcinoma (PDAC). Exclusive application of conventional methods does not appear promising. Deep learning approaches has achieved great success in the computer aided diagnosis, especially in biomedical image segmentation. This paper introduces a framework based on convolutional neural network (CNN) for segmentation of PDAC mass and surrounding vessels in CT images by incorporating powerful classic features, as well. First, a 3D-CNN architecture is used to localize the pancreas region from the whole CT volume using 3D Local Binary Pattern (LBP) map of the original image. Segmentation of PDAC mass is subsequently performed using 2D attention U-Net and Texture Attention U-Net (TAU-Net). TAU-Net is introduced by fusion of dense Scale-Invariant Feature Transform (SIFT) and LBP descriptors into the attention U-Net. An ensemble model is then used to cumulate the advantages of both networks using a 3D-CNN. In addition, to reduce the effects of imbalanced data, a new loss function is proposed as a weighted combination of three classic losses including Generalized Dice Loss (GDL), Weighted Pixel-Wise Cross Entropy loss (WPCE) and boundary loss. Due to insufficient sample size for vessel segmentation, we used the above-mentioned pre-trained networks and fin-tuned them. Experimental results show that the proposed method improves the Dice score for PDAC mass segmentation in portal-venous phase by 7.52% compared to state-of-the-art methods (from 53.08% to 60.6%) in term of DSC. Besides, three dimensional visualization of the tumor and surrounding vessels can facilitate the evaluation of PDAC treatment response.

## 1. Introduction

Pancreatic ductal adenocarcinoma (PDAC), constituting 90% of pancreatic cancers, is one of the main causes of cancer death worldwide (1, 2). At the time of diagnosis, most patients suffer from the advanced stage of disease leading to an overall 5-year survival rate of %8 (3). Only 10–20% of patients have resectable tumors at the time of diagnosis (1). PDAC staging is mainly based on the degree of involvement between the tumor and surrounding vessels such as Superior Mesenteric Artery (SMA) and Superior Mesenteric Vein (SMV). Therefore, 3D visualization of the tumor and adjacent vessels can be useful for determining tumor-vessel involvement and assessing treatment response in pancreas surgery planning (1).

Chemoradiation therapy as a treatment option is suggested for borderline resectable and locally-advanced PDAC. One of the most important stages before each chemoradiation treatment is manual assessment of the tumor volume which is a time-consuming and complex task depending on radiologist’s experience.

Automated volumetric segmentation of critical tumors plays a crucial role in diagnosis and monitoring problems (6). Finding the lesions in organs such as liver, brain and breast is of great interest in medical image segmentation domain (4-6).

CT is the most common modality used for evaluation of pancreatic cancer. Fully-automated segmentation of PDAC in CT scans is indubitably one of the most challenging tumor segmentation tasks (7-9). Main difficulties in automated pancreas and tumor segmentation arise from three aspects: 1) Variability in terms of size, shape and location of pancreas and especially PDAC mass, as illustrated in Fig. 1; 2) the small size of the pancreas and tumor in the whole CT scan; 3) poor contrast around the boundaries (7, 10-11).

**Figure 1.**
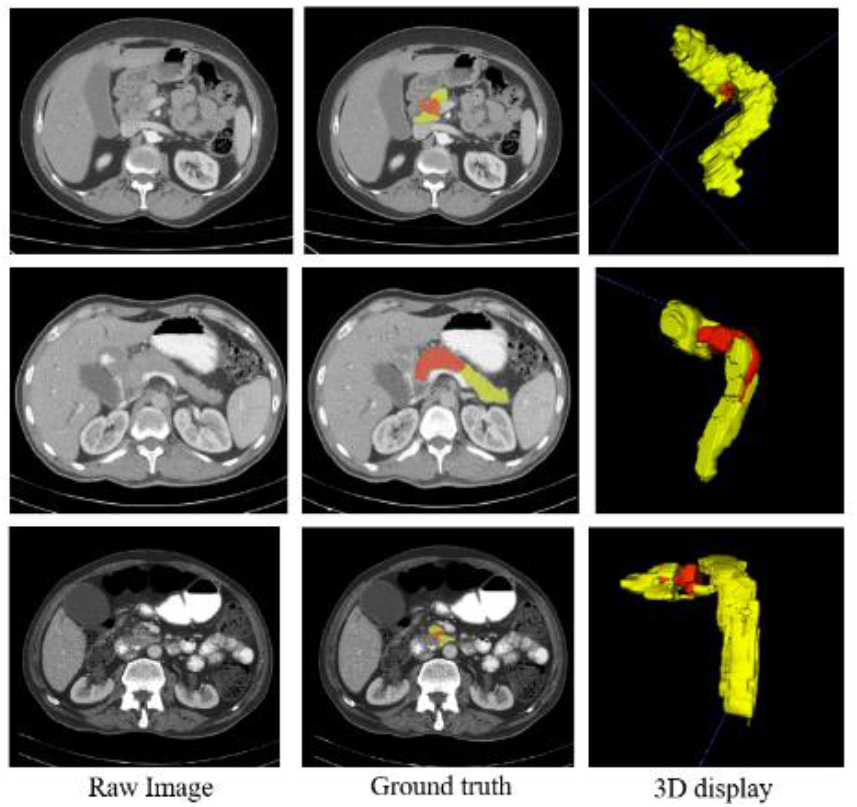
Illustrations of abnormal pancreas, showing the large variations of shape, size and location of PDAC mass. Normal pancreas region is denoted as yellow and PDAC mass is marked as red.

Organ segmentation methods can be generally divided into two categories: Bottom-up and top-down methods. Top-down methods apply a-prior knowledge such as shape models or atlases generated and incorporated into the framework using shape model fitting or image registration (12, 13). Bottom-up approaches use pixel or voxel based labeling or local image similarity grouping (14, 15). The bottom-up methods are more effective in pathological organ segmentation (11).

Recently deep learning has achieved great success in the computer aided diagnosis, especially in the biomedical image segmentation domain (16-19). Several studies focused on the pancreas segmentation using deep learning techniques. In 2018, Oktay et.al (18) proposed self-soft attention mechanism for segmentation of pancreas. Frag et. al (10) applied cascade super pixels for pancreas segmentation on NIH dataset using deep and texture features. Gibson et. al (20) proposed a method based on registration–free deep learning to segment eight organs, including pancreas. Roth et al. (21) introduced an approach based on pre-segmented pancreas framework followed by a refinement convolutional network, which was further improved using holistically-nested network. They proposed AG modules for suppressing the unrelated regions and also highlighting useful regions with salient features simultaneously. Man Y et.al introduced a Deep Q Network (DQN) driven approach with deformable U-Net to segment the pancreas and subsequently achieved state-of-the-art mean DSC 86.93 ± 4.92% on NIH pancreas dataset (22). Zhu et al. (23) proposed a consecutive 3D coarse-to-fine segmentation model, using the bypass structure of ResNet on two datasets, namely NIH pancreas dataset and JHMI pathological pancreas dataset. In another work, Zhou et al. (24) designed a convolutional network model to localize and segment the pancreatic cyst.

To the best of our knowledge, very limited studies have focused on pancreatic tumor segmentation. The reported results show low accuracy for PDAC and other pancreatic tumors. Zhu et. al presented a multi scale coarse to fine segmentation for screening PDAC in CT images, achieving a dice score of 57.3% for PDAC mass segmentation (7). Tureckova et.al (8) proposed a CNN approach using deep supervision and attention gates for segmentation of lesions such as liver tumor and pancreatic tumor. They reached dice score of 54.66% for pancreas-tumor segmentation. Furthermore, Zhang et.al (9) used multi-phase CT images for PDAC segmentation using a large dataset and nnUNet network, which achieved dice scores of 0.709 ± 0.159 and 0.522 ± 0.250 for multi and venous phase respectively. Zhou el.al (10) proposed PDAC segmentation using hyper-pairing network which integrated the information from different phases. They reported dice scores of 63.94 ± 22.74 and 53.08 ± 27.06 using multi and venous phase respectively. The results of these studies indicate the necessity of achieving better performance in PDAC segmentation.

Furthermore, several studies have investigated abdominal blood vessels segmentation in CT images. Farag et al. (25) proposed a method for abdominal vessel segmentation in contrast enhanced CT images. They obtained automated seed points for vessel tracking and generation of statistical models of the vessels using outputs from the multi-organ multi-atlas label fusion and the Frangi vesselness filter. In another work, Oda et al. (26) presented a fully convolutional network (FCN) for abdominal artery segmentation in CT volume. They used a patch-based method for segmentation of the small arteries with an average F1-measure, precision and recall of 87.1%, 85.8%, and 88.4%, respectively.

Texture descriptors such as Scale Invariant Feature Transform (SIFT), Local Binary Pattern (LBP), GLCM and wavelets have shown to be promising in pancreas segmentation and PDAC detection (11, 27-28). Several other studies have fused texture features and deep features to achieve robust segmentation and prognosis (11, 29). Dense SIFT proposed in (30) and Local Binary Pattern (LBP) (31) have been proven to be powerful descriptors providing informative information.

In this study, using a combination of these texture descriptors and convolutional neural networks, a fully automatic system is proposed for segmentation of PDAC mass and surrounding vessels based on improved attention U-Net. The contributions of this work are introduced in a staged framework as follows:

- A localization stage is designed to find the smallest bounding box covering the entire pancreas from the whole CT scan. For this purpose, 3D CNN and 3D LBP are used to extract corresponding sub-volumes from 3D CT scans.
- A finer stage is proposed for automated segmentation of PDAC mass and normal pancreas from cropped sub-volumes in previous stage. For this purpose, Attention U-Net and Texture Attention U-Net (TAU-Net) are integrated with dense SIFT and 3D LBP which provide rich information in spatial domain.
- A hybrid model is developed as an ensemble model to fuse the Attention U-Net and TAU-Net from the fine stage and to achieve better performance in segmentation.
- A new loss function is proposed as a weighted combination of Generalized Dice Loss (GDL), Weighted Pixel-Wise Cross Entropy Loss (WPCEL) and boundary Loss.
- An approach is designed for segmentation of SMA and SMV using fine-tuning of the above-mentioned models.

The reminder of the paper is structured as follows: Section 2 outlines details of the proposed methodology. Experimental results are presented in Section 3. Section 4 are devoted to discussion and conclusion.

## 2. Materials and Methods

In this study, pancreatic ductal adenocarcinoma CT scans from two datasets are used. First dataset is the publicly available and titled Medical Segmentation Decathlon dataset (32). It includes three types of pancreatic tumor (intraductal mucinous neoplasms, pancreatic neuroendocrine tumors, and pancreatic ductal adenocarcinoma). We selected 138 pancreatic ductal adenocarcinoma cases by radiologists in our team. The second dataset was collected at our hospital. It contains 19 retrospectively pathologically proven PDAC cases. Ethics committee of our university had confirmed the study protocol (IR.TUMS.MEDICINE.REC.1397.119). In the self-collected dataset, pancreas, tumor and vessels such as SMA and SMV are labeled in each manually selected slice by two experienced radiologists using ITK-SNAP v.3.6.0 (http://www.itksnap.org/) (33). The resolution of each CT scan in both dataset is 512 × 512 × L, where L is the number of sampling slices along the long axis of body. For data pre-processing, all images are clipped to [0.5, 99.5] percentiles of the intensity value, followed by a min-max normalization on each volume.

### 2.1 Related works

#### 2.1.1 Attention U-Net

U-Net architecture has been revealed to achieve accurate and robust performance in medical image segmentation (19). Attention Gates (AG) were introduced that can be utilized in CNN frameworks for dense label prediction to focus on target structure without additional supervision (17). Attention U-Net, an extension to standard U-Net, applies self-soft attention technique in a feed-forward CNN model for medical image segmentation (18). The sensitivity of the model to target pixels is improved without utilizing complicated heuristics. Using gating module, salient features are highlighted while noisy as well as unrelated responses disambiguate. These operations are performed just before each skip connection to fuse the relevant features. High-level and low-level feature maps are fed into AG module according to Eq. 1.

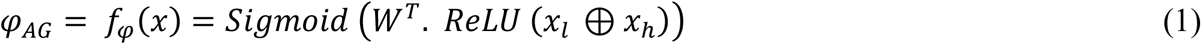

where *φ*_*AG*_ is called the attention coefficient, *W*^*T*^ denote a linear transformation, *x*_*l*_ and *x*_*h*_ represent the low-level and the high-level feature map respectively, *Sigmoid* (*x*) = 1/(1 + *e*^-*x*^), and ⊕ denotes the matrix addition. Attention features are obtained from the element-wise product of the low-level feature map and attention coefficients (Eq. 2).

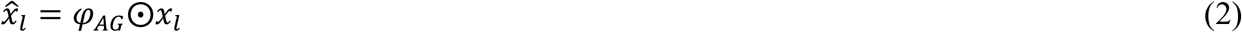

where 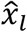 is the attention feature, ⨀ represents element-wise product and *φ*_*AG*_ is the attention coefficient of the higher level features.

These attention features are subsequently concatenated with high level features extracted from the network in decoder path.

#### 2.1.2 Dense SIFT

U-Net and Attention U-Net as classical semantic segmentation networks, are not significantly effective for the challenging pancreatic tumor segmentation task (8). Due to the difficulty of labelling process which leads to a somewhat small sample size, these networks may not benefit from a large number of layers and channels. To solve this issue, we proposed an approach which combines selected traditional salient features with high level features in the decoder path.

One of the most popular feature space transforms proposed by Lowe (34) is SIFT descriptor which includes two stages: key point detector and descriptor. The SIFT algorithm is invariant to rotations, translations and scaling transformations. This property is obtained by characterizing local gradient information around a corresponding detected point of interest.

The original SIFT is a sparse feature representation method for an image while dense feature representation is preferred in pixel classification. Liu et al. (30) proposed the dense SIFT descriptor for object recognition and registration which eliminates the feature detection stage, while preserving the pixel-wise feature descriptor.

#### 2.1.3 2D and 3D Local Binary Pattern (LBP)

We can extend the previous idea of combining classical features with CNN extracted features. Two dimensional LBP, presented by Ojala et al. (35), is an impressive texture descriptor to characterize the local texture patterns through encoding pixel values. It leads to strong capability of rotation and gray level invariance.

Banerjee et al (31) introduced a rotationally invariant three dimensional LBP algorithm where the invariants are constructed implicitly using spherical harmonics for increasing spatial support. A Non-Gaussian statistic measure (kurtosis), was used due to loss of phase information. The number of obtained variables was equivalent to the number of spherical harmonics plus the kurtosis term. This method can locally estimate the invariant features, useful in describing the small patterns.

### 2.2 Segmentation of PDAC mass

The proposed method consists a three-step segmentation process. Due to small occupied portion of CT volumes with desired pancreas region, in the first step, the approximated smallest bounding box for pancreas is localized using proposed 3D LBP sub-volumes and 3D CNNs. Subsequently, the cropped ROI are fed into segmentation network with basis of Attention U-Net to segment pancreas and PDAC mass. We also introduce a modified Attention U-Net, named Texture Attention U-Net (TAU-Net) using information from Dense-SIFT and 3D-LBP. Finally, ensemble approach is applied using a light-weight CNN to take advantage of the mentioned networks. The overall diagram of the proposed method is depicted in Fig. 2.

**Figure 2.**
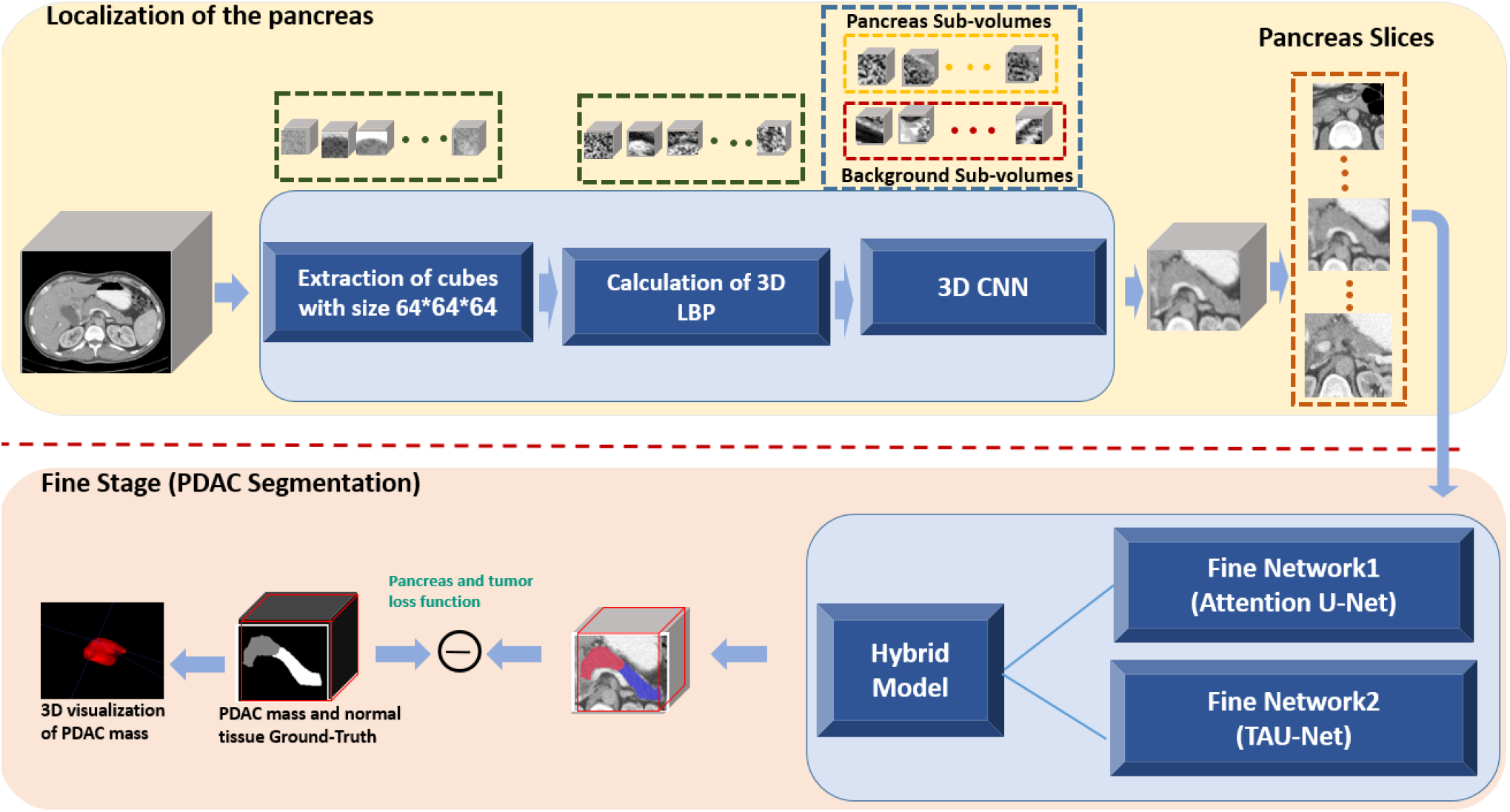
Block Diagram of the proposed segmentation method in the testing phase for segmentation of PDAC

#### 2.2.1 Localization of Pancreas Region

The goal of this stage is to obtain an approximate estimation of where the pancreas volume is located inside the whole 3D CT so as to exclude irrelevant organs and tissues. Although 2D CNN frameworks have achieved considerable progress, contextual information cannot be preserved along z-axis. On the other hand, training 3D networks on a large number of sub-volumes needs considerable computational power and memory. Inadequate number of samples, on the contrary, leads to inappropriate results.

As a solution for this problem, we propose a new approach based on LBP texture descriptor. Instead of training the network using original sub-volumes, informative sub-volumes are extracted utilizing 3D-LBP to be fed to the network. As pancreas region occupy a small portion of the CT volume, sub-volumes with size of 64×64×64 pixels are randomly sampled in the training phase. The extracted sub-volumes either cover a fraction of the pancreas/tumor voxels (foreground) or contain background regions. Although the background volume is much bigger than the foreground in the original image, we selected balanced sets of sub-volumes for the training phase. In the testing phase, a sliding window was applied to the entire CT volume with spatial stride of 50 along both *X* and *Y* axes, and the stride of 20 along the *Z* axis. Subsequently, 3D-LBP are also calculated for each volume, with the same size as the sub-volumes with parallel counterpart pixels. Such 3D-LBP volumes are then fed into a 3D-CNN, composed of five convolution layers with max-pooling, batch normalization (BN) (36) and leaky rectified linear unit (ReLU) as nonlinear activation function. In each convolution block, two 3 × 3 × 3 convolution kernels with stride 1 and no padding are used to reduce the network parameters. The pooling layers are applied on a 2*×*2*×*2 sliding window with stride 2, padded with 2 pixels. The network is trained by Adam optimizer with 5 mini-batchs and a base learning rate of 0.01 benefitting from a polynomial decay (gamma = 0.1) in a total of 30 iterations. Training process is carried out using cross entropy loss. Figure 3 summarizes the proposed localization process of the pancreas within the whole CT volume.

**Figure 3.**
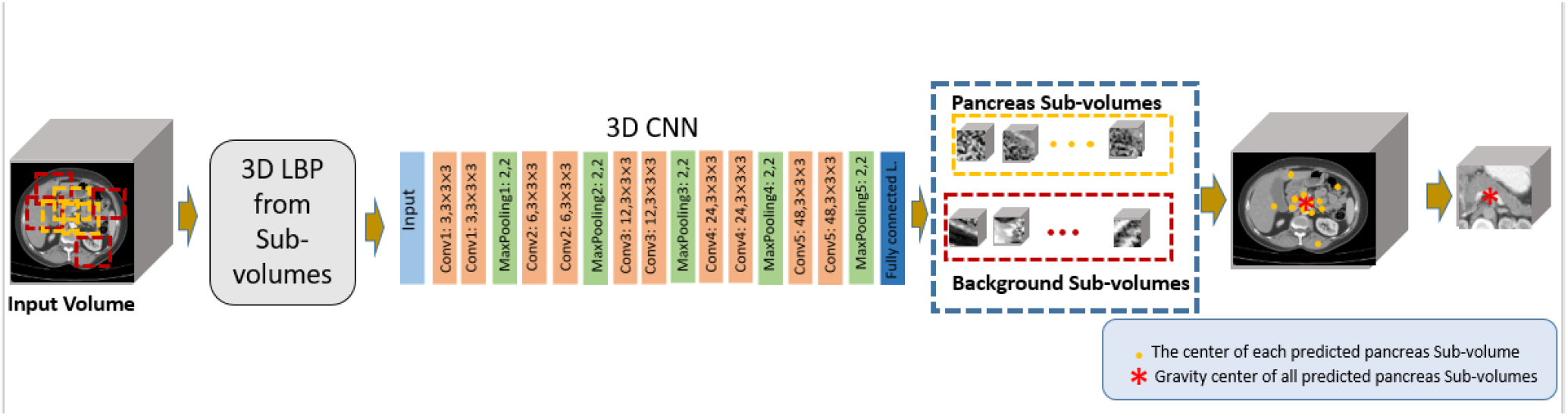
Block diagram of the proposed method for localization of the pancreas region

In the testing phase, to find the location of pancreas, the center of each predicted pancreas sub-volume is obtained and the gravity center of these points are subsequently calculated as the center of detected pancreas volume. A bounding box is finally placed around the center to cover the whole pancreas.

#### 2.2.2 Fine Stage (Segmentation of PDAC mass)

Using previous stage, we obtained smallest sub-volume from the whole CT scan which can be had the benefit of the dimensionality reduction to facilitate the fine segmentation. It’s worth mentioning that the fine segmentation results are mapped back to their places in the entire CT scan using the locations of the rectangular cube that determines where to cut bounding box from the whole CT scan.

##### Texture Attention U-Net (TAU-Net)

In the fine stage, extracted 3D sub-volume for each case is cut into a series of 2D slices to be fed into the next new network. The proposed network in this stage is attention U-Net integrated with texture descriptors. To detail, dense SIFT and 3D LBP are used to enrich the deep features extracted in convolutional layers and design the TAU-Net as shown in Fig 4 (a). TAU-Net has an architecture similar to attention U-Net. The input images are progressively down sampled with five convolution steps which extracts higher level image representations *x*_*h*_. Each step consists a convolution block (two 3 × 3 convolution layers followed by batch normalization and ReLU activation). A max-pooling operation subsequently is used with a kernel size 2 × 2 and stride of 2 to reduce the size of the transitional feature maps. At the decoder path we used deconvolution layers, AGs and Texture Attention Gates (TAGs) along with skip connections. AGs and TAGs are applied for skip connection to aggregate information from multiple scales. In addition to the generated deep features from the previous network layers, low-level features, such as dense SIFT and 3D-LBP, are used to provide a more comprehensive representation of the pathological characteristics of the tissues. Texture Attention Block (TA Block) is responsible for integration of these efficient features with features from the decoding path.

**Figure 4.**
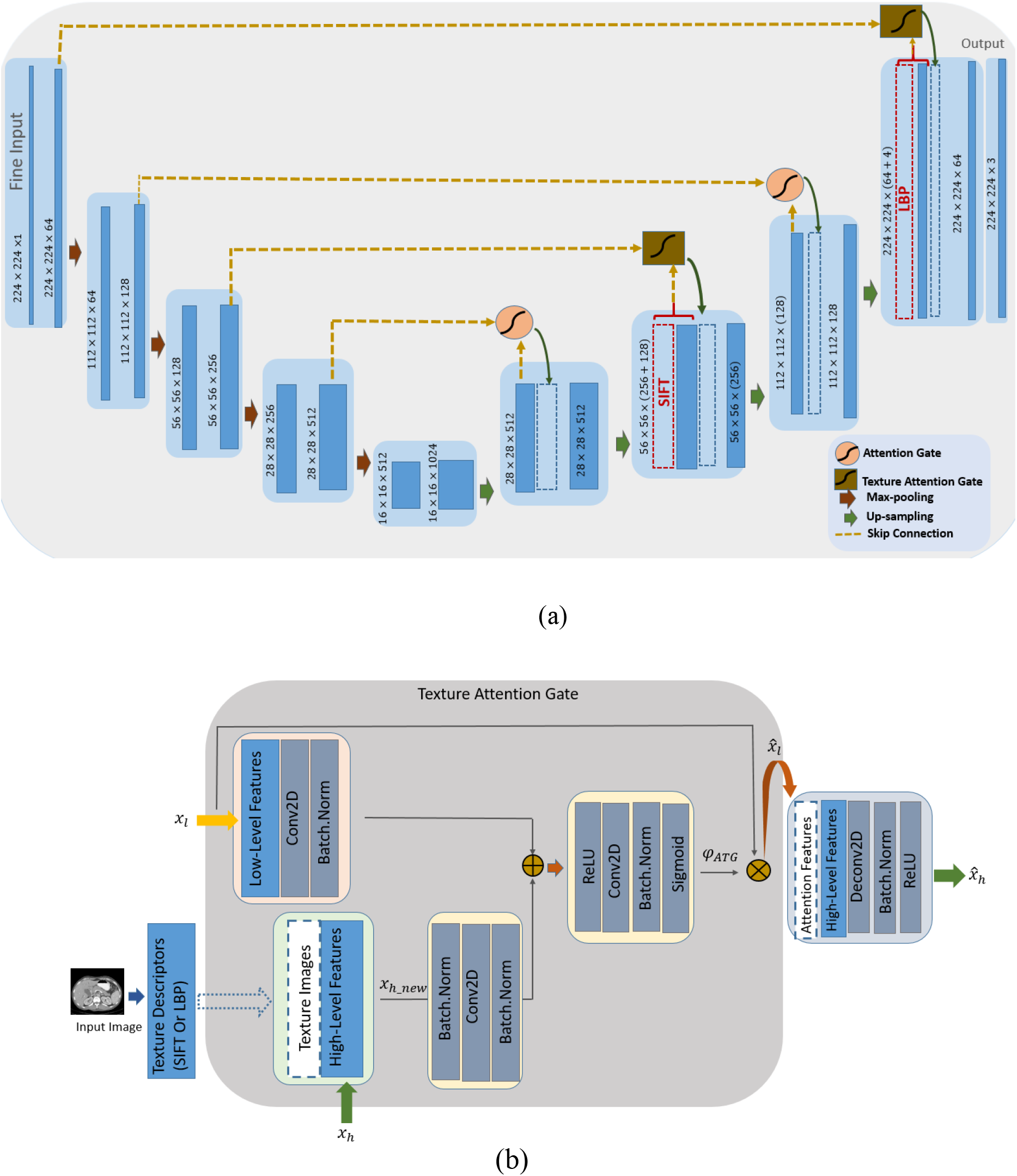
(a) TAU-Net architecture, (b) Block diagram of the proposed TABlock.

##### Texture Attention Block (TA Block)

In the TA Block (Fig. 4(b)), high-level feature maps (*x*_*h*_) are first concatenated with texture features (TXFea) to result in informative features. The output of this stage (*x*_*h*_*new*_) is one of the inputs to the gate module. Newly produced high-level feature maps (*x*_*h*_*new*_) and also low-level feature maps (*x*_*l*_) from the coding stage are combined to compose the inputs of the attention gate (Eq.3).

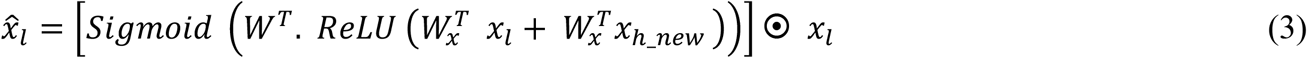

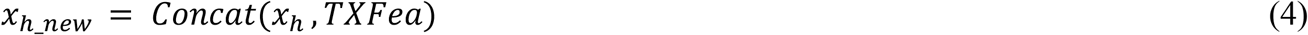

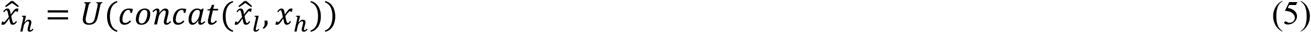

where, 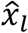 represents attention features, *x*_*h*_*new*_ is high-level features concatenated with texture features, 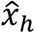 denotes the input to the next stage in the decoder path and U denotes up-sampling.

The Best configuration is finally achieved emprically based on the network performance. As can be seen from Fig. 4, SIFT features are integrated into the network at the second up-sampling layer. Since the size of the feature map in this layer is a quarter of the input image size, SIFT features are calculated with a sampling size of four. Besides, LBP was integrated into network at the last layer which has the same size of the input image. The network is trained using the Adam optimizer in 10 mini-batches and a base learning rate of 5×10e-5 with polynomial decay (gamma = 0.1). We have also used random rotation, flip and shift to augment the data. A random 70-30 split was used for training-testing phases.

#### 2.2.3 Proposed hybrid model

In this study, Attention U-Net and TAU-Net are used for segmentation of pancreas and the tumor using CT images. To benefit the advantages of both networks, a novel but simple approach based on hybrid models (37, 38) is proposed using a 3D-CNN to aggregate the outputs of these networks. First, three predicted masks, namely pancreas, tumor and background are generated using each of the two segmentation networks leading to six predicted masks as the inputs to the 3D-CNN aggregator network. This network consists of one convolution layer and a softmax function as shown in Fig. 5. The 3D convolution layer predicts label of each pixel using its neighbors along three axes.

**Figure 5.**
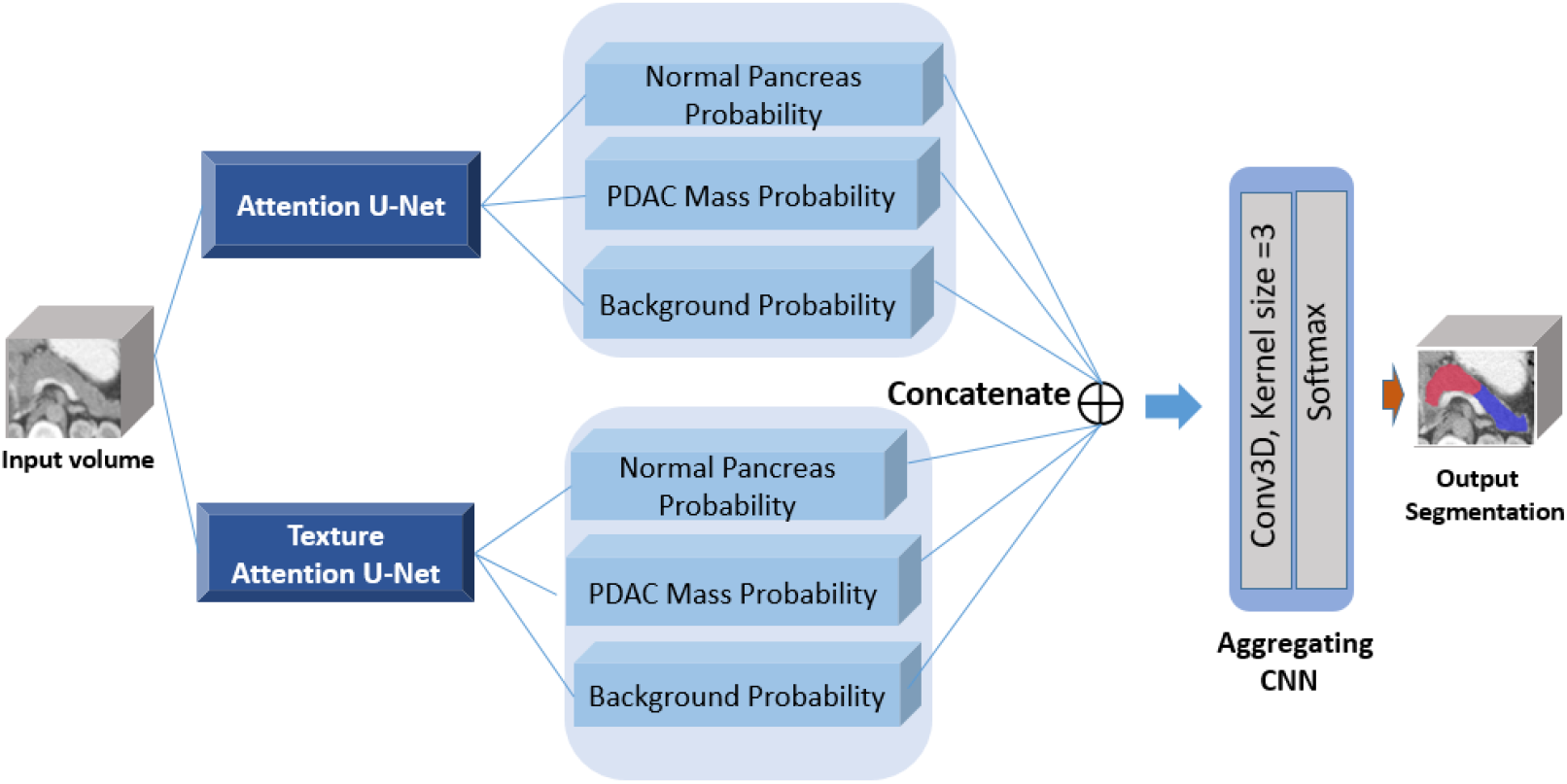
Block diagram of the proposed hybrid model

#### 2.2.4 Loss Function

Standard losses such as cross entropy loss and Dice loss cannot handle the imbalanced tasks appropriately (39). Hence, to deal with small foreground as well as large intra class variation of region sizes, we use three different loss functions, namely GDL, boundary loss and WPCE.

##### Weighted Pixel-Wise Cross Entropy (WPCE)

WPCE loss is Defined as follow:

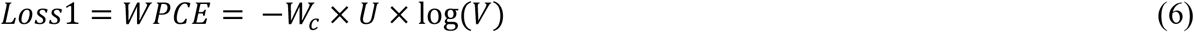

where *U* represents ground truth and *V* is the predicted binary mask. In this study, the sample weights for each class was selected to be the ratio of total number of background pixels *n*_*B*_ and total number of each class (pancreas or tumor) pixels *n*_*c*_, 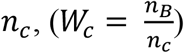.

##### Generalized Dice Loss (GDL)

GDL is defined in eq. (7) (40).

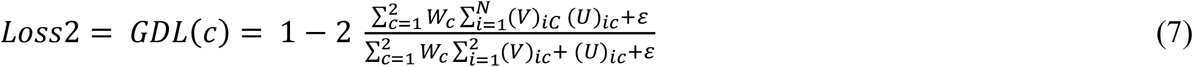

where 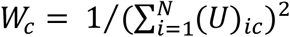 is responsible for being the invariant to properties of the different label sets.

##### Boundary Loss

Because of low contrast around the boundary of pancreas and tumor, exact determination of tumor and healthy tissue boundary can be challenging. To diminish this issue, we also use the differentiable version of the boundary loss proposed in (41). The boundaries of the ground truth and predicted segments are obtained using max-pooling operation as follow:

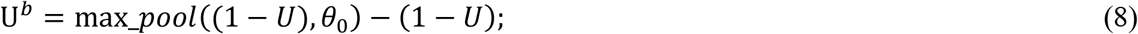

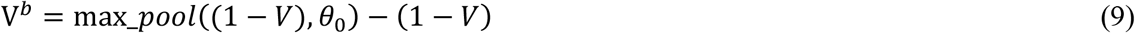

where a pixel-wise max-pooling operation is applied to obtain the inverted ground truth and predicted binary segments with a kernel size *θ*_0_. Then an extended boundary is constructed using Euclidean distances of the pixels from boundaries:

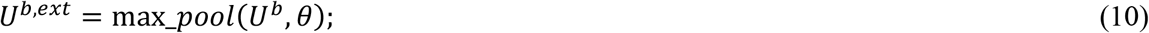

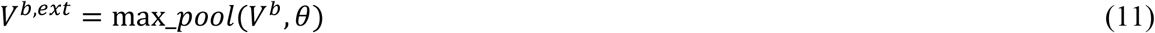

Subsequently, *P* (precision) and *R* (recall) is calculated using the above constructed maps. Boundary metrics (*BF*_1_) (42) can be calculated using precision and recall. Finally, the loss function is defined as:

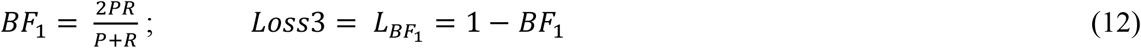

Ultimately, the weighted sum of the above loss functions is used as follow:

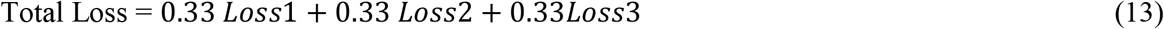

### 2.3 Segmentation of vessels

PDAC staging is mainly based on the degree of involvement between the tumor and surrounding vessels. So segmentation of surrounding vessels of the pancreas can be a crucial task in pancreas surgery planning. As mentioned at the beginning of section two, 19 self-collected 3D CT scans have vessel labels. This sample size is insufficient to train the vessel segmentation networks from scratch. In order to overcome this limitation, we utilized the models trained for PDAC segmentation, namely Attention U-Net, TAU-Net and hybrid models, as pre-trained models and fine-tuned them for vessel segmentation. The data is augmented by translation, flipping and rotation of each image to make the dataset larger 12 times. The initial learning rate of the Adam optimizer is 1e-5. We train the networks for 30 epochs.

### 2.4 Evaluation Metrics

The following metrics were used to evaluate the pancreas and PDAC mass segmentation: Dice similarity coefficient (DSC), Recall, Precision.

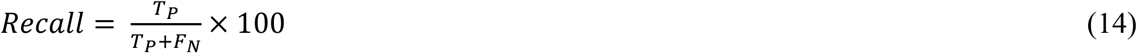

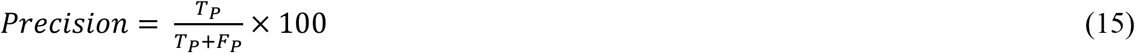

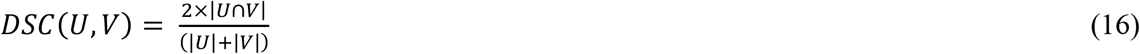

where *U* and *V* denote ground-truth and predicted binary masks respectively.

All algorithms are implemented using the publicly available PyTorch package (43). Training of all models was carried out using a machine with a single NVIDIA GFORCE GTX1080Ti with 11GB memory.

## 3. Results

### 3.1 Pancreas localization

The 3D CNN is trained to localize the pancreas in the whole CT scan. Table 1 shows the results of training this network with original images and 3D LBP sub-volumes using different number of 3 × 3 filters.

**Table 1.** Pancreas classification results for 64×64×64 sub-volumes using 3D CNN

| Input Data | No. of Filters | Sensitivity | Specificity | Accuracy |
| --- | --- | --- | --- | --- |
| Original sub-volume | 2 | 0.47 | 0.93 | 0.86 |
| Original sub-volume | 3 | 0.52 | 0.94 | 0.87 |
| 3D LBP sub-volume | 2 | 0.75 | 0.95 | 0.93 |
| **3D LBP sub-volume** | **3** | **0.95** | **0.94** | **0.95** |

### 3.2 PDAC mass segmentation

The Attention U-Net and TAU-Net were trained using images obtained from the previous stages. As shown in figure 4a, the best performance of TAU-Net was obtained when SIFT images were added into the second layer and the LBP images into the last layer of the decoder path. The results of Attention U-Net, TAU-Net and hybrid networks with manual and automated localization of pancreas region are reported in Table 2.

**Table 2.**
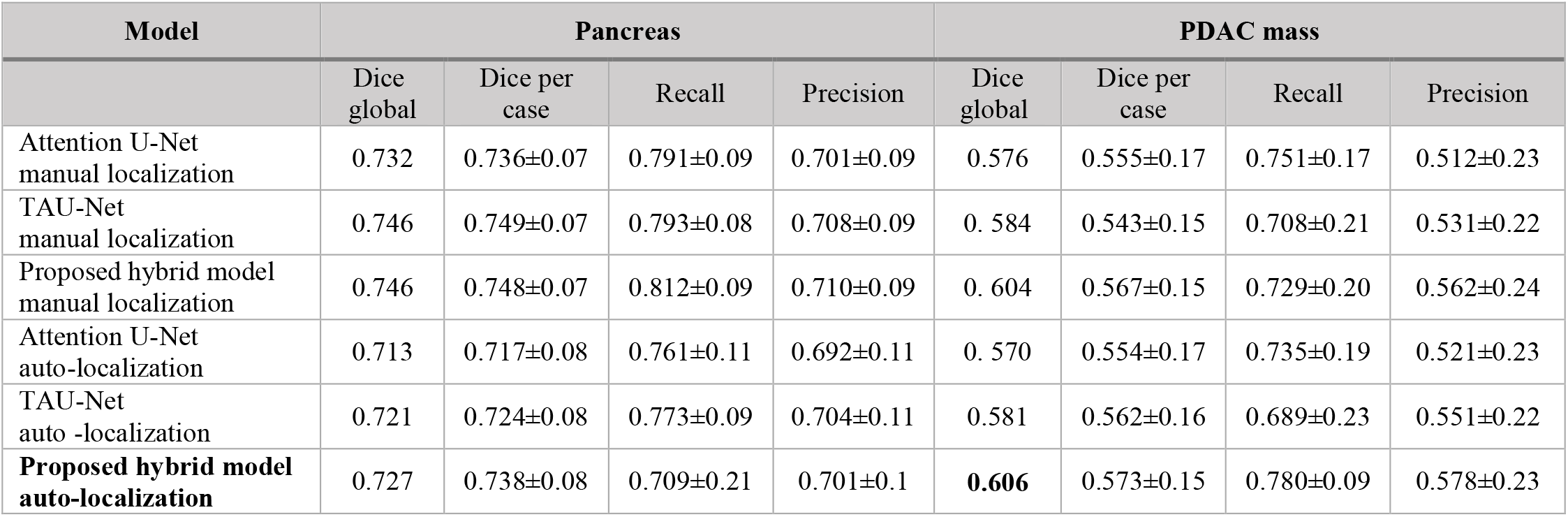
Pancreas and PDAC mass segmentation results using attention U-Net, TAU-Net and hybrid model.

| Model | Pancreas |  |  |  | PDAC mass |  |  |  |
| --- | --- | --- | --- | --- | --- | --- | --- | --- |
|  | Dice global | Dice per case | Recall | Precision | Dice global | Dice per case | Recall | Precision |
| Attention U-Net manual localization | 0.732 | 0.736±0.07 | 0.791±0.09 | 0.701±0.09 | 0.576 | 0.555±0.17 | 0.751±0.17 | 0.512±0.23 |
| TAU-Net manual localization | 0.746 | 0.749±0.07 | 0.793±0.08 | 0.708±0.09 | 0.584 | 0.543±0.15 | 0.708±0.21 | 0.531±0.22 |
| Proposed hybrid model manual localization | 0.746 | 0.748±0.07 | 0.812±0.09 | 0.710±0.09 | 0.604 | 0.567±0.15 | 0.729±0.20 | 0.562±0.24 |
| Attention U-Net auto-localization | 0.713 | 0.717±0.08 | 0.761±0.11 | 0.692±0.11 | 0.570 | 0.554±0.17 | 0.735±0.19 | 0.521±0.23 |
| TAU-Net auto-localization | 0.721 | 0.724±0.08 | 0.773±0.09 | 0.704±0.11 | 0.581 | 0.562±0.16 | 0.689±0.23 | 0.551±0.22 |
| **Proposed hybrid model auto-localization** | 0.727 | 0.738±0.08 | 0.709±0.21 | 0.701±0.1 | **0.606** | 0.573±0.15 | 0.780±0.09 | 0.578±0.23 |

As it can be seen from Table 2, the hybrid model with *“*ensemble*“* approach using a 3D CNN shows the best performance in PDAC mass segmentation. To show the performance of the localization stage, in addition to the results of fully automated pancreas and PDAC mass segmentation, we have also reported the results of feeding manual localization of the pancreas region to the attention U-Net, TAU-Net and hybrid network in Table 2. Furthermore, figure 6 shows the qualitative segmentation results for different CT slices where rows 1 and 2 depict two samples from MSD dataset while row 3 and 4 represent results for two samples from self-collected dataset. 3D reconstruction of the results using ITK-SNAP (Fig. 6 (d)) shows an anatomical and realistic view with very small irregularities.

**Figure 6.**
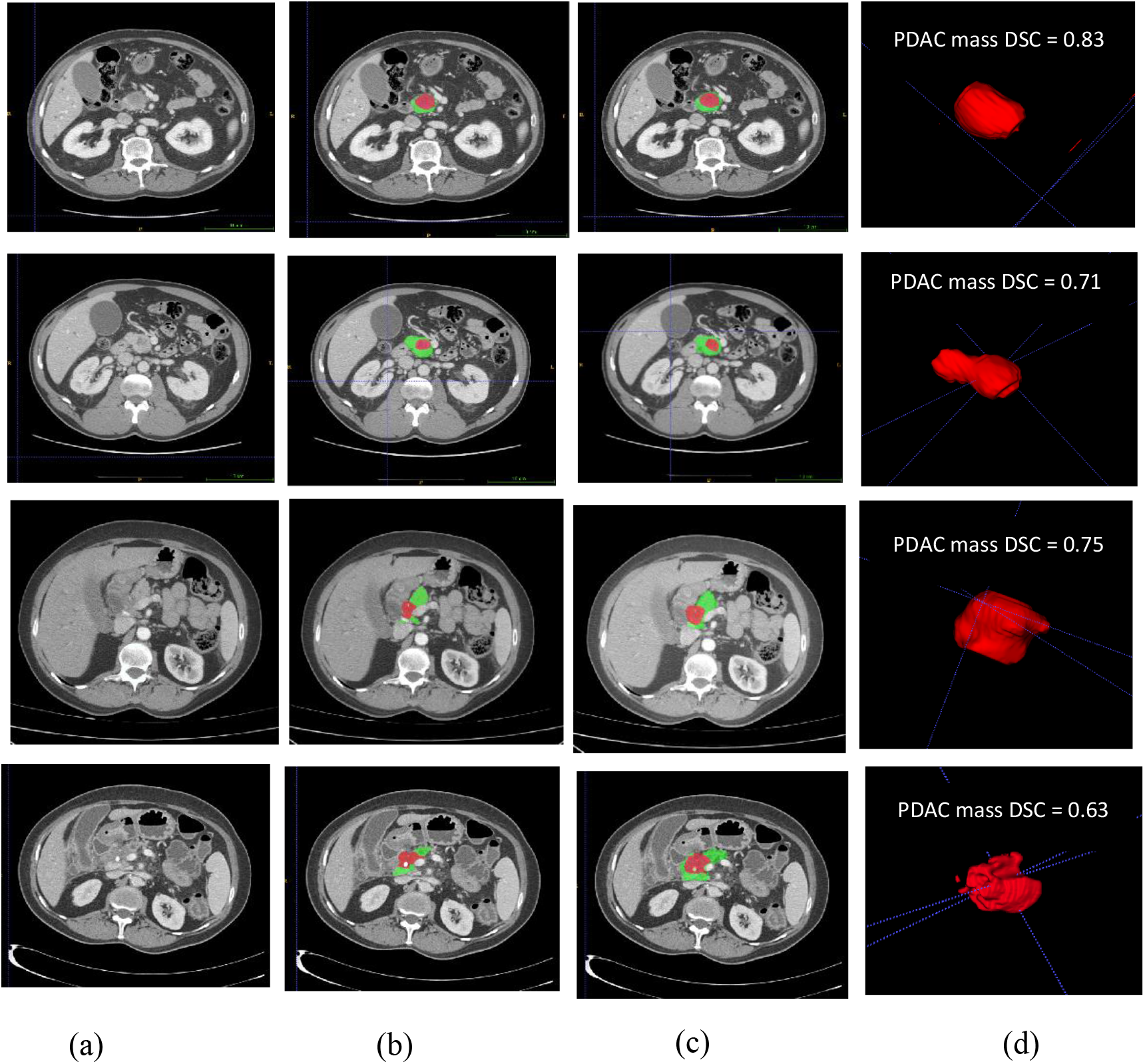
Final segmentation results of the pancreas and PDAC mass using the hybrid network, a) Original image, b) Ground truth (red indicates PDAC mass and green presents normal pancreatic tissue), c) Final segmentation, d) 3D visualization of PDAC mass.

### 3.3 Vessel Segmentation

We have compared the performance of three networks, namely the pre-trained Attention U-Net, TAU-Net and hybrid model for segmentation of vessels. Segmentation results for two samples are reported in fig. 7. For further validation purposes, DSC, recall and precision for all networks are reported in Table 3. Furthermore, fig. 7 (d and i) shows the segmented PDAC mass and vessels. Three visualization of PDAC mass and surrounding vessels is presented in figure 7 (e and l).

**Table 3.** Vessel segmentation results using pre-trained Attention U-Net, TAU-Net and hybrid model

| Model | Superior Mesenteric Artery (SMA)<br>Segmentation |  |  | Superior Mesenteric Vein (SMV)<br>Segmentation |  |  |
| --- | --- | --- | --- | --- | --- | --- |
|  | Dice | Recall | Precision | Dice | Recall | Precision |
| Pre-trained Attention U-Net | $0.76 \pm 0.08$ | $0.79 \pm 0.12$ | $0.76 \pm 0.09$ | $0.68 \pm 0.02$ | $0.77 \pm 0.06$ | $0.62 \pm 0.07$ |
| Pre-trained TAU-Net | $0.80 \pm 0.06$ | $0.88 \pm 0.07$ | $0.74 \pm 0.09$ | $0.70 \pm 0.09$ | $0.66 \pm 0.12$ | $0.76 \pm 0.06$ |
| **Pre-trained hybrid model** | $0.81 \pm 0.06$ | $0.87 \pm 0.09$ | $0.76 \pm 0.07$ | $0.73 \pm 0.04$ | $0.81 \pm 0.10$ | $0.68 \pm 0.03$ |

**Figure 7.**
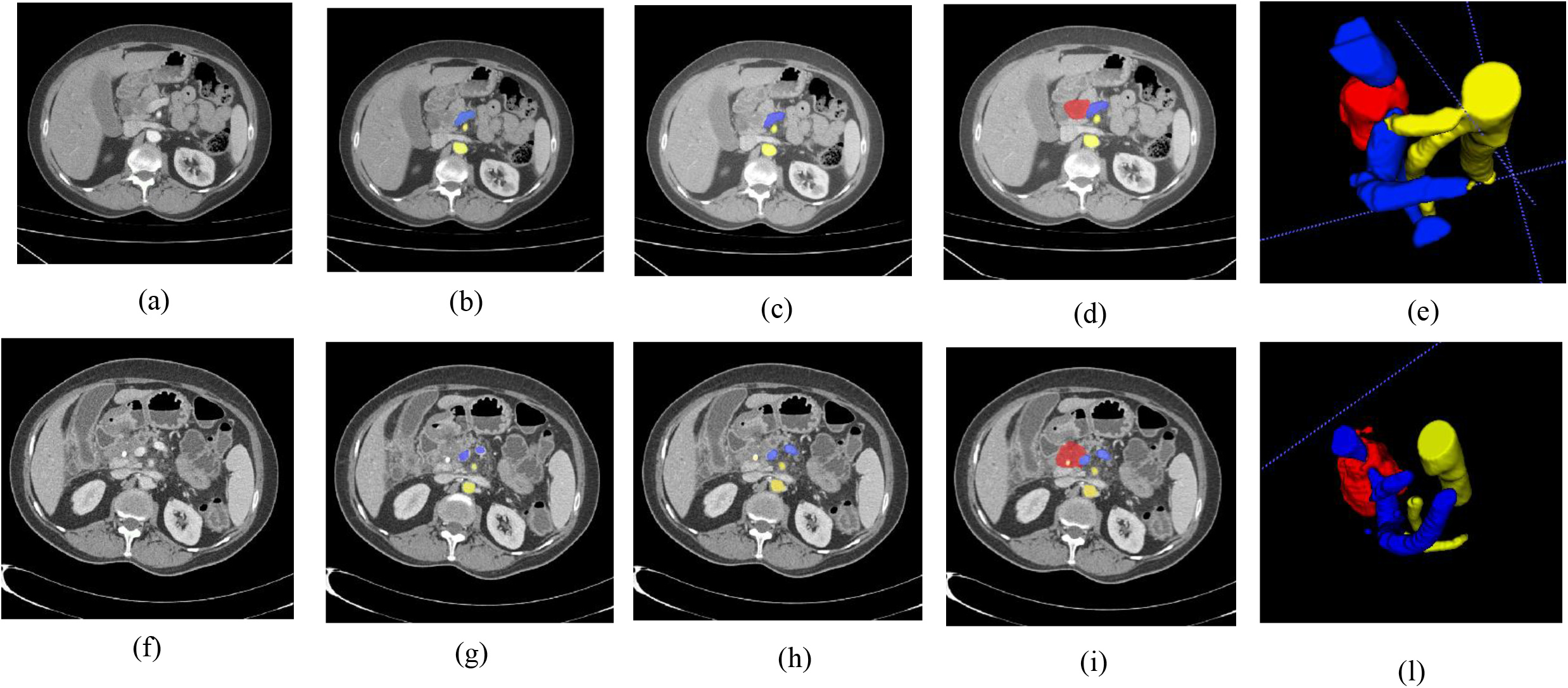
Final segmentation and 3D visualization of the PDAC mass and surrounding vessels. a) Original image for a case before chemoradiation therapy, b) Ground truth for SMA (yellow) and SMV (blue), c) Segmentation results of the SMA and SMV using pre-trained hybrid model, d) Final segmentation of PDAC mass, SMA and SMV, e) 3D visualization of PDAC mass, SMA and SMV. f) Original image for the same case after chemoradiation therapy, g) Ground truth for SMA (yellow) and SMV (blue), h) Segmentation results of the SMA and SMV using pre-trained hybrid model, i) Final segmentation of PDAC mass, SMA and SMV, l) 3D visualization of PDAC mass, SMA and SMV.

## 4. Discussion and Conclusion

In this paper, we introduced a fully automatic convolutional neural network for segmentation of PDAC mass and abdominal vessels such as SMA and SMV in CT images. We also introduced TAU-Net as a modified Attention U-Net, using information from dense SIFT and 3D-LBP. As PDAC staging is mainly based on the degree of involvement between the tumor and surrounding vessels, segmentation of PDAC mass as well as surrounding vessels can be an important task, which is already addressed in this study.

To localize the pancreas slices in the whole CT image, although a single network could possibly be used in both cascaded stages (coarse and fine steps), but a desired outcome can only be expected from extremely complicated. To overcome this challenge, the proposed method used a simple 3D CNN with inputs from the 3D LBP maps extracted from sub-volumes of the original images. With this approach, instead of applying a large number of original images, a small sample size of informative 3D LBP maps resulted in an accuracy of 95% in classification between pancreas and background sub-volumes. Furthermore, as it can be found according to Table 2, by feeding manual and automated localization of the pancreas region to the segmentation stage networks, automated and manual localizations of pancreas region show comparable performances.

Although the performance of the segmentation stage might improve with the integration of contextual information through 3D architecture networks (7, 44), due to the limited size of the dataset and the involved high computational cost, 2D architectures are selected in this work to analyse each CT slice individually. The disadvantage of the 2D architecture is that each slice does not include all of the contextual and geometric information due to the absence of the disregarded z-axis information.

To overcome this weakness partially, the hybrid model using a simple 3D CNN was introduced to incorporate information from the adjacent slices to the segmentation of the intended slice. One may assume that the superior performance of the hybrid network compared to each network alone, is due to using data from adjacent slices as well as utilizing a combination of texture features and deep features.

To incorporate texture information, we examined fusion of SIFT, 2D-LBP and slices of 3D-LBP into the selected layers of the network. Experimental results showed that 3D-LBP performs better than the 2D-LBP as expected. Figure 6 compares 2D-LBP and 3D-LBP of a single slice. The 2D slice of the 3D-LBP is much more informative than both the original image and the 2D LBP.

**Figure 6.**
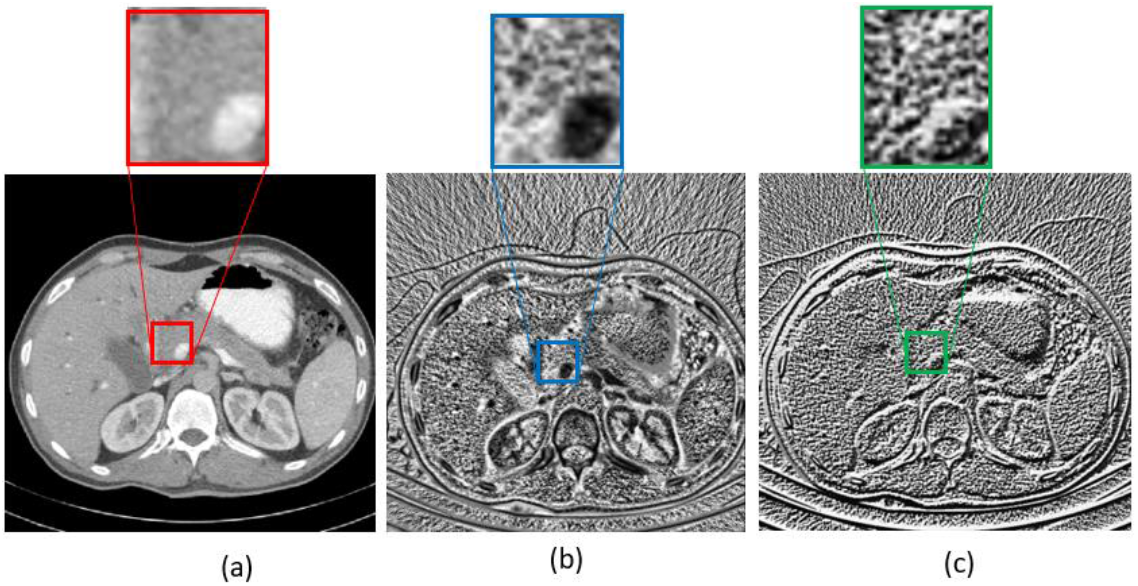
(a) Original image (a slice of the original volume), b) LBP slice obtained from the 3D LBP of the original volume, c) 2D LBP obtained from the original image.

The best result for fusion of the 3D LBP into the network obtained when it was added to the last layer in the decoder path. SIFT maps were also fused into the network. The SIFT maps are calculated using different step sizes and each obtained map is concatenated into its matched layer in the decoding path. This leads to mutual advantages of high and low level SIFT features during the decoding. The mean dice value for PDAC mass segmentation using (1) Attention U-Net + LBP, (2) Attention U-Net + SIFT_1 + LBP, (3) Attention U-Net + SIFT_2 + LBP, (4) Attention U-Net + SIFT_4 + LBP and (5) Attention U-Net + SIFT_8 + LBP were 0.54, 0.55, 0.53, 0.58 and 0.55 respectively. SIFT_*n*, means that the SIFT map with step size of *n* is added to the corresponding layer in the decoder path. The best performance among all architectures belong to Attention U-Net with SIFT_4 + LBP. It seems that in detection of the targeted texture, neither very low level nor extreme high level SIFT features provide enough information.

To reduce the effects of unbalanced data (pancreas vs background) we examined a tailored set of loss functions consisting of weighted combinations of three classic losses namely GDL, WPCE and boundary loss. WPCE shows the best performance compared to the other two losses. But the best final results have been achieved using the balanced combination of all mentioned losses. It seems that the best results are obtained with this combination due to usage of high and low resolution losses simultaneously.

Previous pancreatic tumor segmentation approaches have led to Dice scores between 0.52 to 0.71, depending on tumor types (PDAC and others), utilizing single or multi CT phase (non-contrast, arterial, venous), dataset size and image quality. But compared to studies which use portal venous phase data, current imaging protocol and routine, the proposed method benefits from a significant improvement of up to 7.52% (from 53.08 (10) to 60.6). One of the most remarkable superiorities of our study over other previous works is segmentation and 3D visualization of PDAC mass and surrounding vessels which can facilitate the assessment of treatment response in pancreas surgery planning.

In vessel segmentation we had a small sample size (19 self-collected case). Therefore, pre-trained networks, obtained from PDAC segmentation, are fine-tuned for this task and the results are reported in Table 3. As it can be seen, TAU-Net shows the better performance than the Attention U-Net on the segmentation of vessels and PDAC mass segmentation respectively. It can be concluded that this network can be promising in segmentation of medical images with small sample size.

As the future step, we propose fusion of 3D texture descriptors into the 3D networks. In addition, we suggest using GLCM and multi-resolution features such as dual tree complex wavelet transform. Furthermore, in the hybrid model, fine tuning of the final combined network may be a good solution.

By conducting this study, we tried to enhance the diagnosis and visualization results by using current imaging protocols and routines. Another successive phase may contain manipulating the protocols and even the media for achieving better results. Though this work mainly focuses on segmentation of PDAC mass, we can apply the proposed idea to the segmentation of another pancreas tumor such as intraductal mucinous neoplasms and pancreatic neuroendocrine tumors as well as other lesions like liver, brain and lung. Three dimensional visualization of PDAC mass and surrounding vessels can facilitate the assessment of treatment response in pancreas surgery planning. However, in order to be integrated into clinical applications, 3D visualization software is a must in further developments.

